# Linking immunity and nutrition: Vital DNA extracellular trap formation in *Dictyostelium discoideum* reveals an ancestral strategy for microbial management

**DOI:** 10.64898/2026.03.20.713133

**Authors:** Antonia Ramos-Guzmán, Paulina Aguilera-Cortés, Sebastián Farías, Ian Pérez, Viviana Barros, Boris Riveros, Tabata Soto, Mauricio Hernández, Camilo Berríos-Pastén, Diego Rojas, Andrés E. Marcoleta, Francisco P. Chávez

**Affiliations:** Laboratorio de Microbiología de Sistemas, Departamento de Biología, Facultad de Ciencias, Universidad de Chile, Santiago, Chile; Grupo de Microbiología Integrativa, Laboratorio de Biología Estructural y Molecular, Departamento de Biología, Facultad de Ciencias, Universidad de Chile, Santiago, Chile; Departamento de Biotecnología, Universidad Tecnológica Metropolitana (UTEM), Santiago, Chile; Melisa Institute, Chile

## Abstract

Extracellular traps (ETs) were originally described in neutrophils as DNA-based structures that immobilize microbes and contribute to innate immunity. Subsequent studies revealed that ET formation occurs across diverse immune cell types and can proceed through non-lytic mechanisms involving mitochondrial DNA release. Whether ETosis also operates outside classical immune contexts and what its ancestral functions may be remain incompletely understood. Here, we show that vegetative, phagocytic cells of the social amoeba *Dictyostelium discoideum*, a professional bacterial predator with phagocytic mechanisms conserved with those of mammalian innate immune cells, deploy extracellular DNA traps in response to bacterial cues. ET formation is selectively induced by specific lipopolysaccharide variants, is not triggered by canonical neutrophil NET inducers, and occurs through a vital ETosis mechanism that preserves membrane integrity and feeding capacity. Ultrastructural analyses provide the first visualization of extracellular traps in Amoebozoa, revealing extracellular filamentous networks that physically capture bacteria. Molecular characterization demonstrates that amoeboid ETs are enriched in mitochondrial DNA and harbor a dynamic proteomic repertoire dominated by mitochondrial components, DNA-associated proteins, and multiple antibacterial effectors. Notably, ET composition varies with the bacterial stimulus, indicating that ETs are not static structures but rather responsive extracellular assemblies. Together, these findings establish ET formation as a regulated response in a unicellular phagocyte and suggest that extracellular traps may have originally functioned in microbial management during feeding, prior to their elaboration as immune effectors in multicellular organisms.

**Highlights:**

- Vegetative *Dictyostelium discoideum* cells deploy mitochondrial DNA-based extracellular traps in response to bacterial cues.
- ET formation occurs through a vital, non-lytic mechanism that preserves membrane integrity and feeding capacity.
- Extracellular traps exhibit stimulus-dependent composition and are enriched in mitochondrial and antimicrobial proteins.
- ETosis functions as an ancestral strategy for microbial containment and management beyond canonical immune contexts.

## Introduction

Microbial life has evolved for billions of years within complex ecological networks dominated by competition, predation, and nutrient limitation. In these environments, bacteria have interacted extensively with diverse eukaryotic hosts, including protists and plants, long before the emergence of animals. Accumulating evidence suggests that many traits now associated with virulence, along with mechanisms underlying antimicrobial resistance, reflect ancestral biological functions shaped by prolonged ecological interactions rather than adaptations to modern pathogenic lifestyles ^1,2^. Protozoan predation has been proposed as a significant evolutionary driver selecting for bacterial traits that enhance survival against phagocytic eukaryotic cells, many of which later function as virulence determinants in animal hosts ^3,4^.

Extracellular traps (ETs) represent a compelling example of how ancient ecological functions may have been repurposed during the evolution of innate immunity. In vertebrates, neutrophil extracellular traps (NETs) are DNA-based structures composed of chromatin decorated with antimicrobial proteins that immobilize and restrict microbes. Since their discovery in neutrophils ^5,6^, ETosis has been increasingly recognized as a conserved antimicrobial strategy across multiple specialized immune cell types ^7^. Early models initially described ETosis as a suicidal process involving nuclear DNA release; however, subsequent studies revealed alternative, non-lytic forms of ETosis. Notably, neutrophils and eosinophils can release mitochondrial DNA (mtDNA) to form extracellular traps while remaining viable, demonstrating that ET formation does not necessarily entail cell death ^8,9^. While ETs play important roles in host defense, excessive or dysregulated ET production contributes to inflammatory and autoimmune pathologies, underscoring the need to understand both their regulation and evolutionary origins.

Despite extensive investigation in vertebrate immune systems, the ancestral functions and evolutionary emergence of ETs remain poorly understood. Protists provide an ideal framework to explore these questions, as feeding and defense are intrinsically coupled in unicellular predators. The social amoeba *Dictyostelium discoideum* is a soil-dwelling, bacterivorous eukaryote that relies on phagocytosis for nutrition and survival ^10^. Remarkably, *D. discoideum* exhibits cellular behaviors analogous to those of professional immune cells, including chemotaxis, phagocytosis, phagosome maturation, and efficient intracellular killing of microbes ^11,12^. At the molecular level, *D. discoideum* encodes homologs of several innate immune components, including Toll/Interleukin-1 receptor (TIR)-like proteins, NOD-like receptors (NLRs), and the NADPH oxidase NoxA, which enables reactive oxygen species (ROS) production comparable to the respiratory burst observed in neutrophils ^13,14^. Bacterial ingestion by amoebae further involves compartmentalized phagosomal trafficking and regulated delivery of hydrolytic enzymes, processes that closely parallel those of mammalian phagocytes ^15–17^.

During nutrient deprivation, *D. discoideum* transitions from unicellular growth to a multicellular developmental program, forming migrating slugs composed of differentiated cell types. Within this multicellular context, specialized sentinel (S) cells have been shown to deploy mtDNA-based extracellular traps in response to lipopolysaccharide (LPS), contributing to antimicrobial defense within the multicellular slug during development ^18^. However, whether vegetative and phagocytic cells, which are the predominant cell types during growth and feeding, retain the capacity to form ETs remains unclear. This distinction is fundamental, as vegetative cells continuously interact with bacteria not only as potential pathogens but also as nutritional resources, raising the possibility that ET formation in this context may serve functions beyond those associated with specialized immune cells.

In this study, we investigate whether vegetative *D. discoideum* cells retain the capacity to form DNA-based extracellular traps in response to bacterial cues, and whether ET formation in this context differs from that described in specialized immune cells. We examine how ET deployment is regulated, its molecular composition, and its cellular origin in vegetative amoebae. We further explore the functional relevance of ET formation during interactions with both pathogenic and edible bacteria, addressing whether extracellular traps may contribute to microbial control in the context of feeding. Finally, we assess whether ET formation in vegetative amoebae occurs through non-lytic mechanisms, drawing parallels with mitochondrial DNA-based ETosis described in other organisms.

By integrating cellular, molecular, and functional analyses, this work aims to determine whether ETosis in *D. discoideum* represents a broader cellular strategy in which phagocytic feeding is coupled to mitochondrial DNA release and extracellular control of bacteria, and to establish this organism as a model for investigating the evolutionary origins of extracellular trap biology.

## Results

### Vegetative *D. discoideum* cells release extracellular DNA structures in response to LPS

Previous studies demonstrated the capacity of *D. discoideum* sentinel (S) cells to form mitochondrial DNA-based extracellular traps (ETs) in response to *Klebsiella pneumoniae* cells and its lipopolysaccharide (LPS) ^18^. However, whether phagocytic vegetative cells can also release ETs remained unexplored. To address this question, we developed a fluorescence-based assay to quantify extracellular DNA release using Sytox Green, a membrane-impermeant dye that selectively stains extracellular DNA.

Vegetative cells of *D. discoideum* (AX2 and AX4 strains) were exposed to increasing concentrations of LPS from *K. pneumoniae* (2.5, 5, and 10 µg/mL), and Sytox fluorescence was monitored kinetically using a microplate reader. A robust dose-dependent increase in fluorescence was observed over time (Figure 1A), with the highest signal detected at 10 µg/mL after 4 hours of stimulation (Figure 1B). Control samples displayed only minimal fluorescence during this time window, with a gradual increase at later time points likely reflecting physiological stress under starvation conditions. Based on these kinetics, the 4-hour time point was selected for subsequent analyses.

**Figure 1.**
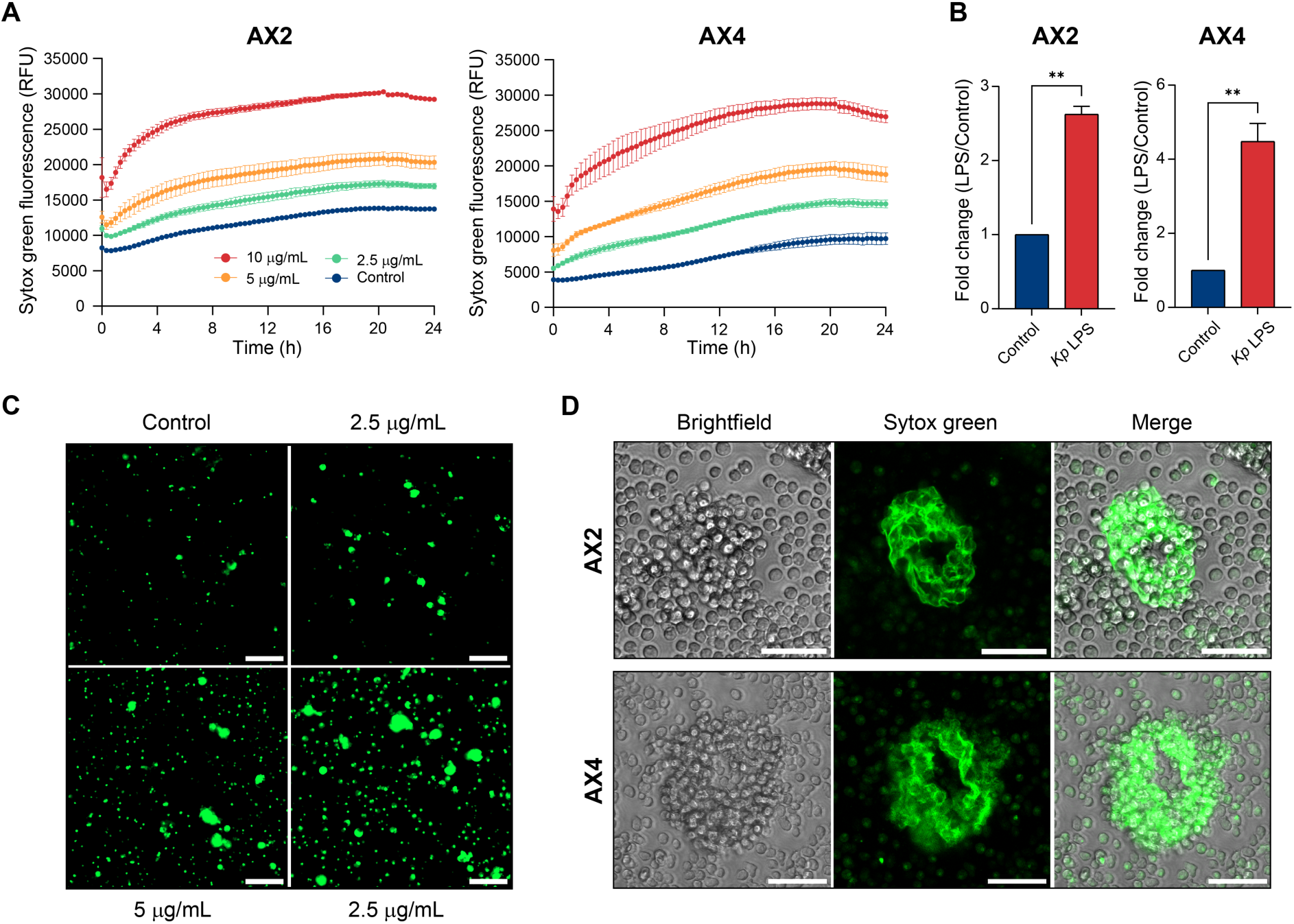
Dose-dependent induction of extracellular traps (ETs) in vegetative *Dictyostelium discoideum*. (A) *D. discoideum* AX2 and AX4 cells were stimulated with increasing concentrations of *K. pneumoniae* LPS (*Kp* LPS; 2.5, 5, and 10 µg/mL) or buffer as a control. ET production was quantified by measuring Sytox Green fluorescence over time. (B) At 4 h post-stimulation, the fold change in fluorescence intensity relative to the untreated control indicates a significant increase in ET formation in both strains at the high LPS concentration (10 µg/mL). Data are presented as mean ± SD (n = 3 biological replicates). Statistical significance was assessed using the Welch’s t-test (**: *p* < 0.01). (C) Fluorescence microscopy showing DNA-positive ET structures stained with Sytox Green in the presence of increasing concentrations of *K. pneumoniae* LPS. Scale bar: 50 µm. (D) Aggregative clusters surrounded by extracellular DNA in both AX2 and AX4 cells. Scale bar: 50 µm.

Fluorescence microscopy confirmed the presence of extracellular DNA structures consistent with ETs in LPS-treated cells (Figure 1C), whereas control samples displayed only marginal fluorescence after prolonged incubation, likely reflecting physiological stress under starvation conditions. Confocal microscopy revealed prominent extracellular trap-like networks, occurring primarily at clusters formed by several cells (Figure 1D; Movie S1). Most cells retained a rounded morphology across all conditions, suggesting that membrane integrity was preserved. To verify that the Sytox signal reflected extracellular DNA, LPS-stimulated cells were treated with Turbo DNase. DNase treatment resulted in a pronounced loss of extracellular Sytox fluorescence under microscopy, whereas intracellular fluorescence remained largely unaffected. We also observed a marked reduction in Sytox fluorescence in both the AX2 and AX4 strains, as measured by the microplate reader assay (Figure 2B). Consistently, agarose gel electrophoresis of extracellular nucleic acids revealed a DNA smear that was abolished upon DNase treatment (Figure 2C). Together, these results demonstrate that vegetative *D. discoideum* cells release extracellular DNA structures in response to LPS in a dose-dependent manner, extending previous observations made in sentinel cells to the vegetative growth stage.

**Figure 2.**
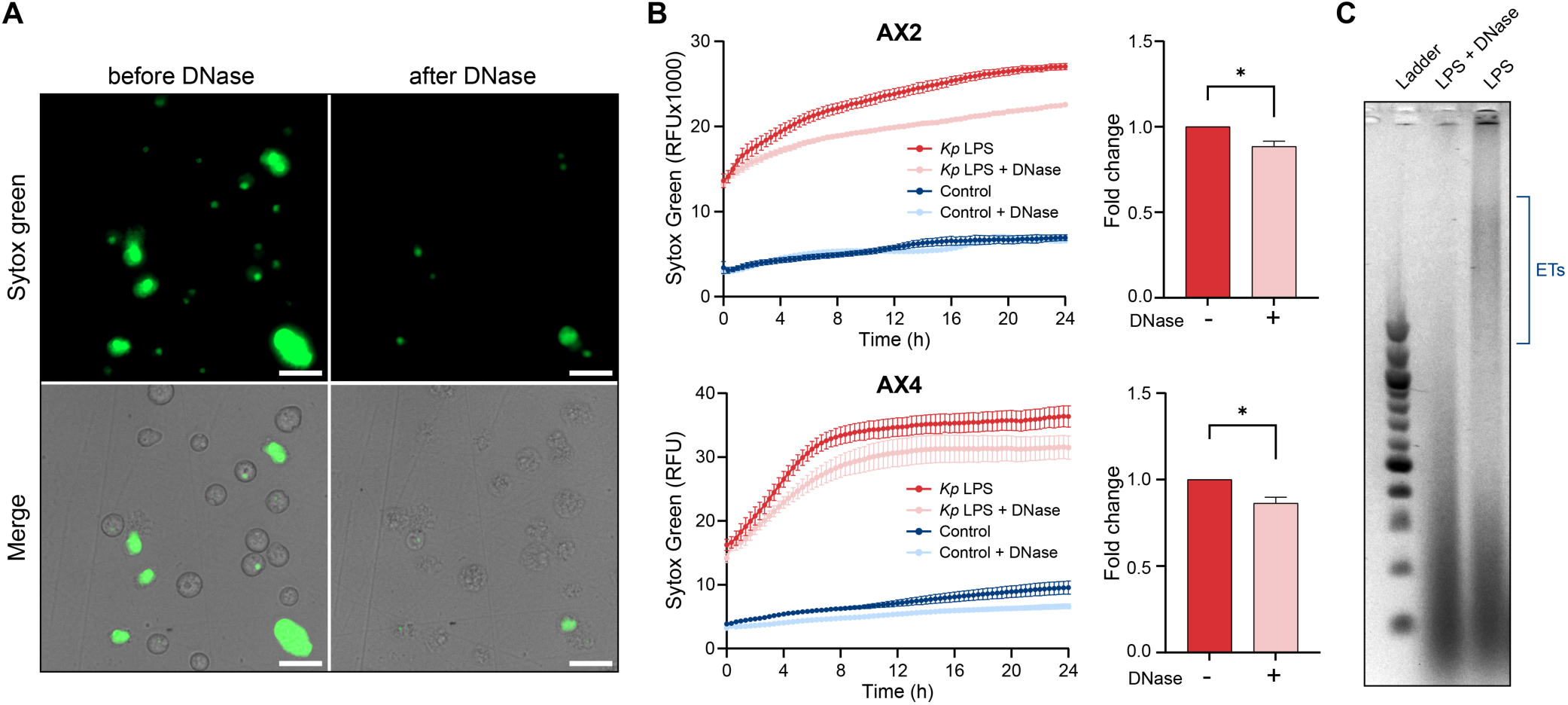
DNase confirms the DNA-based nature of ET structures. (A) Microscopy showing loss of extracellular Sytox Green signal after DNase treatment. Scale bar: 20 µm. (B) Sytox Green fluorescence kinetics with and without DNase treatment in LPS-stimulated (10 µg/mL) AX2 and AX4 cells. At 4 h post-stimulation, the fold change in fluorescence intensity relative to the untreated control indicates a significant decrease in ET formation after DNase treatment in both strains. Data are presented as mean ± SD (n = 3 biological replicates). Statistical significance was assessed using Welch’s t-test (*: p < 0.05). (C) Agarose gel electrophoresis showing DNA degradation in the ET fraction after DNase treatment.

### ET formation in vegetative *D. discoideum* is selectively induced by distinct bacterial LPS variants

To examine whether ET formation varies with the bacterial origin of LPS, we compared responses to LPS from *K. pneumoniae*, *Escherichia coli*, and *Pseudomonas aeruginosa*. ET formation differed among LPS sources. LPS from *K. pneumoniae* induced the strongest response, whereas *E. coli* LPS elicited a weak and variable effect. In contrast, *P. aeruginosa* LPS showed intermediate activity, with a clear dose-dependent increase in fluorescence between 2.5 and 10 µg/mL (Figure S1A). Quantification at 4 hours revealed pronounced source-dependent differences (Figure S1B), and fluorescence microscopy confirmed the presence of Sytox-positive extracellular structures that increased with LPS concentration (Figure S1C).

### Vital ETosis in vegetative *D. discoideum*: DNA ET formation without loss of viability

Having established that ET formation in vegetative *D. discoideum* is selectively induced by specific bacterial LPS variants, we next asked whether this response reflects a cytotoxic process or a regulated, non-lytic form of ETosis. A defining feature distinguishing classical neutrophil NETosis from alternative forms of extracellular trap (ET) formation in other cell types is the relationship between DNA release and cell death. While NETosis in neutrophils is often associated with lytic cell death, non-lytic forms of ETosis have been described, in which viable cells actively extrude DNA while preserving membrane integrity and cellular functions, most notably through the release of mitochondrial DNA ^9^.

To assess the impact of LPS-induced ET formation on cellular viability, we first employed a plaque-based phagocytosis assay that measures the ability of amoebae to survive, proliferate, and feed on bacteria. AX2 and AX4 cells were exposed to 10 µg/mL *K. pneumoniae* LPS for 6 hours and subsequently co-plated with edible *K. pneumoniae* KpGe on SM agar. After 72 hours, LPS-treated amoebae formed numerous phagocytic plaques comparable to those of untreated controls (Figure 3A), indicating preserved viability and feeding competence. In contrast, treatment with cytotoxic agents such G418 resulted in a marked reduction in plaque formation, validating the sensitivity of the assay.

**Figure 3.**
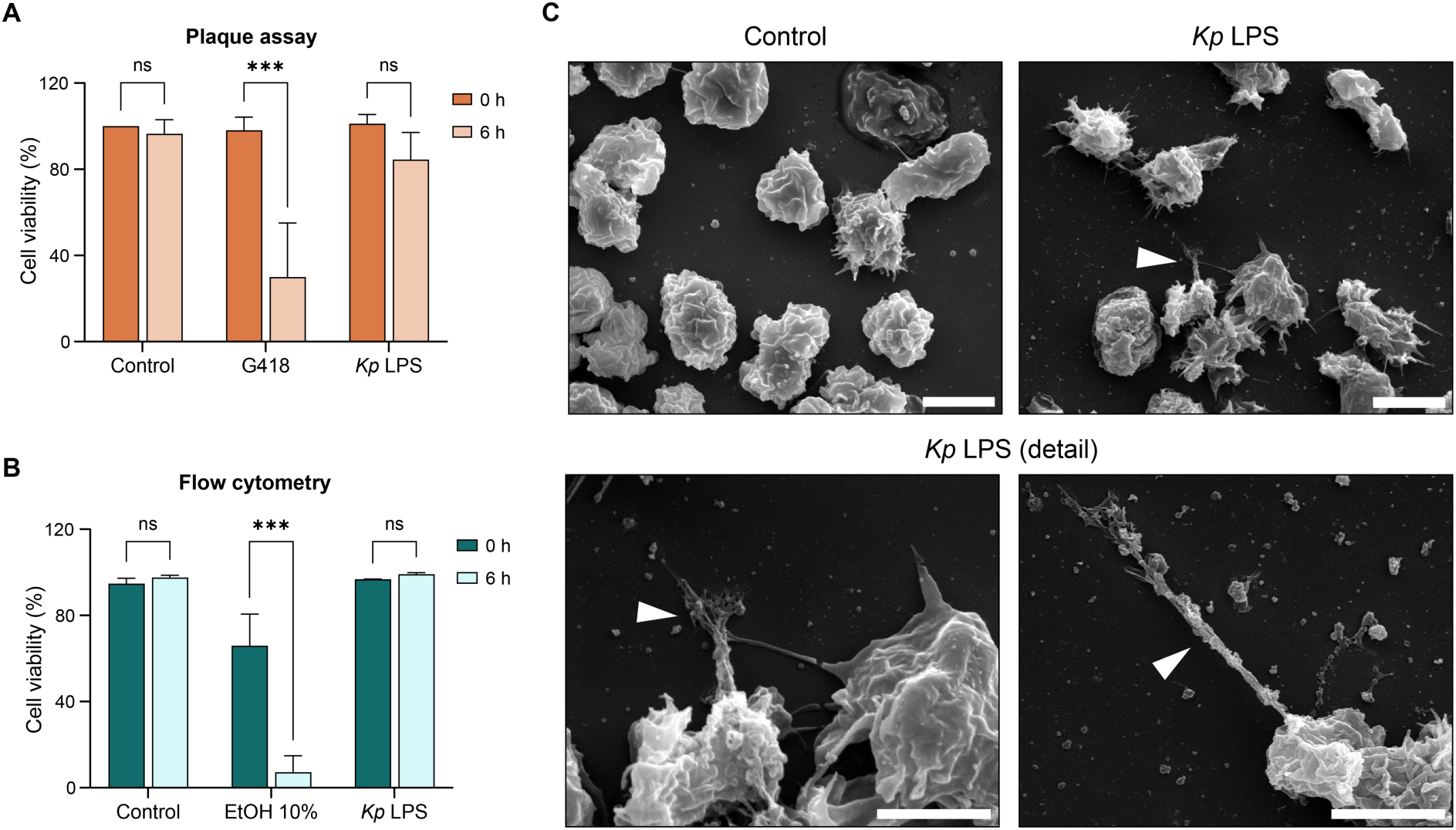
ET formation does not compromise cell viability. (A) AX2 cells viability assay by plaque formation after treatment with *K. pneumoniae* LPS or G418 at 0 h and 6 h. (B) AX2 cells viability assay by flow cytometry using propidium iodide, after treatment with *K. pneumoniae* LPS or ethanol at 0 h and 6 h. Data are presented as mean ± SD (n = 3 biological replicates). Statistical significance was assessed using two-way ANOVA with Bonferroni’s multiple comparisons test (***: p < 0.001). (C) ET ultrastructure visualization by SEM 4 h post-stimulation with *K. pneumoniae* LPS. Arrowheads indicate the presence of fibrillar structures projected from *D. discoideum* cells, consistent with extracellular DNA fibers. Scale bar: 10 µm (upper panels); 5 µm (lower panels).

To independently assess membrane integrity, AX2 cells were analyzed by flow cytometry after propidium iodide (PI) staining, which selectively labels cells with compromised plasma membranes. After 6 hours of LPS exposure, PI incorporation remained negligible and indistinguishable from untreated controls (Figure 3B). In contrast, exposure to ethanol (10%) induced a pronounced increase in PI-positive cells, consistent with extensive membrane damage and cell death. Together, these results demonstrate that LPS-induced ET formation in vegetative *D. discoideum* does not compromise membrane integrity or overall cell viability.

Structural insights into ET formation were further obtained using scanning electron microscopy (SEM). Under control conditions, vegetative amoebae displayed a rounded morphology with a relatively smooth surface (Figure 3C). In contrast, LPS-stimulated cells exhibited pronounced extracellular filamentous structures extending from the cell surface, forming web-like networks that connected neighboring cells and surrounding material. High-magnification SEM images revealed thin fibrillar projections consistent with extracellular DNA strands, while cell bodies retained intact morphology without signs of membrane rupture or collapse. These ultrastructural features are consistent with active ET deployment and, to our knowledge, represent the first SEM-based ultrastructural visualization of ET-like structures in Amebozoa.

Together, functional viability assays, membrane integrity measurements, and ultrastructural analyses demonstrate that ET formation in vegetative *D. discoideum* occurs through a non-cytotoxic mechanism, consistent with a vital form of ETosis.

### Classical ET Inducers and alternative PAMPs do not trigger ET formation in *D. discoideum*

Having established that LPS from *K. pneumoniae* robustly induces ET formation in vegetative *D. discoideum*, we next examined whether other classical inducers of ETosis or bacterial pathogen-associated molecular patterns (PAMPs) elicit similar responses. In neutrophils and other vertebrate immune cells, phorbol 12-myristate 13-acetate (PMA) is a potent activator of NADPH oxidase-dependent ETosis, while flagellin, a TLR5 agonist, activates innate immune signaling pathways ^19–21^.

To test whether *D. discoideum* responds to these stimuli, AX2 cells were exposed to increasing concentrations of PMA ( 50, 500, and 1000 nM), and ET formation was measured by Sytox Green fluorescence. None of the PMA concentrations tested elicited a significant increase in fluorescence relative to untreated controls (Figure S2A). We next evaluated the response to bacterial flagellin at concentrations of 100, 200, and 300 ng/mL. Under these conditions, Sytox Green fluorescence remained comparable to control levels, with no detectable increase in extracellular DNA release (Figure S2B).

Together, these results demonstrate that classical ET inducers and alternative bacterial PAMPs such as PMA and flagellin do not trigger ET formation in vegetative *D. discoideum* under the conditions tested.

### Mitochondrial and antimicrobial signatures define DNA extracellular traps

To define the molecular composition of DNA-based extracellular traps (ETs) formed by vegetative *D. discoideum* cells, we established a fractionation workflow coupled to quantitative PCR and proteomic analyses. Following stimulation with *K. pneumoniae* LPS, cultures were separated into three fractions: the secretome (total supernatant), the ET fraction (supernatant enriched in material adjacent to adhered cells), and the cell pellet (Figure 4A). The ET fraction was treated with DNase to release DNA-associated proteins, which were subsequently analyzed by qPCR and LC-MS/MS-based quantitative proteomics.

**Figure 4.**
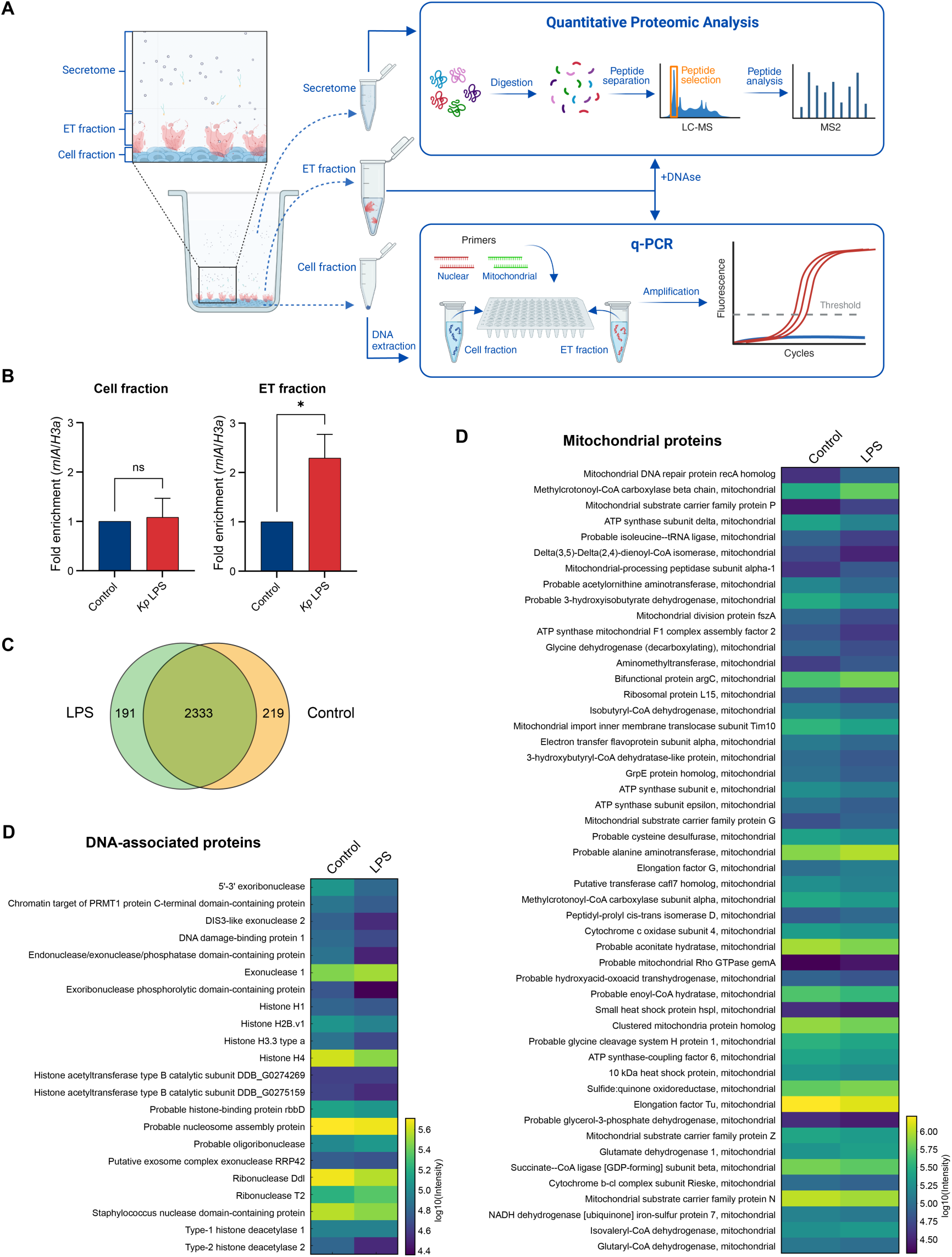
Mitochondrial DNA and proteins dominate ET composition. (A) Schematic representation of sample collection (secretome, ET, and cell fractions) and processing prior to qPCR and proteomic analyses. (B) qPCR analysis showing fold enrichment of mitochondrial (rnlA) and nuclear (H3a) genes in ET versus cell fractions. Data are presented as mean ± SD (n = 3 biological replicates). Statistical significance was assessed using Welch’s t-test (*: p < 0.05). (C–D) Proteomic comparison of control and LPS-induced ETs, including overlap and mitochondrial content. Proteomic profiling of DNase-treated ET fractions revealed a complex and reproducible protein repertoire under both control and LPS-induced conditions. Comparison of protein identities revealed substantial overlap across conditions, with only a limited subset of proteins unique to either state (Figure 4C). Across both conditions, mitochondrial proteins were strongly enriched, including ATP synthase subunits, tricarboxylic acid (TCA) cycle enzymes, mitochondrial ribosomal proteins, and components of oxidative metabolism (Figure 4C, right), supporting a conserved mitochondrial contribution to ET-associated material.

To determine the origin of extracellular DNA, we first quantified mitochondrial and nuclear DNA using qPCR targeting the mitochondrial gene *rnlA* and the nuclear gene *H3a*. LPS stimulation resulted in a significant enrichment of mitochondrial DNA (mtDNA) in the ET fraction, whereas nuclear DNA levels remained unchanged (Figure 4B). No corresponding enrichment was observed in the cell pellet. These results indicate that extracellular DNA released during ET formation is selectively enriched in mtDNA, consistent with an active, stimulus-dependent process.

In addition, numerous DNA-associated proteins were detected, including histones (H2A, H2B, H3, and H4), ribonucleases, exonucleases, helicases, and chromatin-modifying enzymes (Figure 4C, lower left), reinforcing the DNA-centric nature of these extracellular structures.

Beyond mitochondrial and DNA-associated components, ETs were also enriched in proteins with known or predicted antimicrobial activity (Figure 5A). These included phospholipases, cysteine proteases, nucleases, and glycosidases, several of which have been previously implicated in bacterial killing, digestion, or virulence attenuation in *D. discoideum*. Notably, multiple antibacterial proteins identified in ETs correspond to factors recently characterized as contributors to antimicrobial defense and bacterial clearance in amoebae ^22,23^.

**Figure 5.**
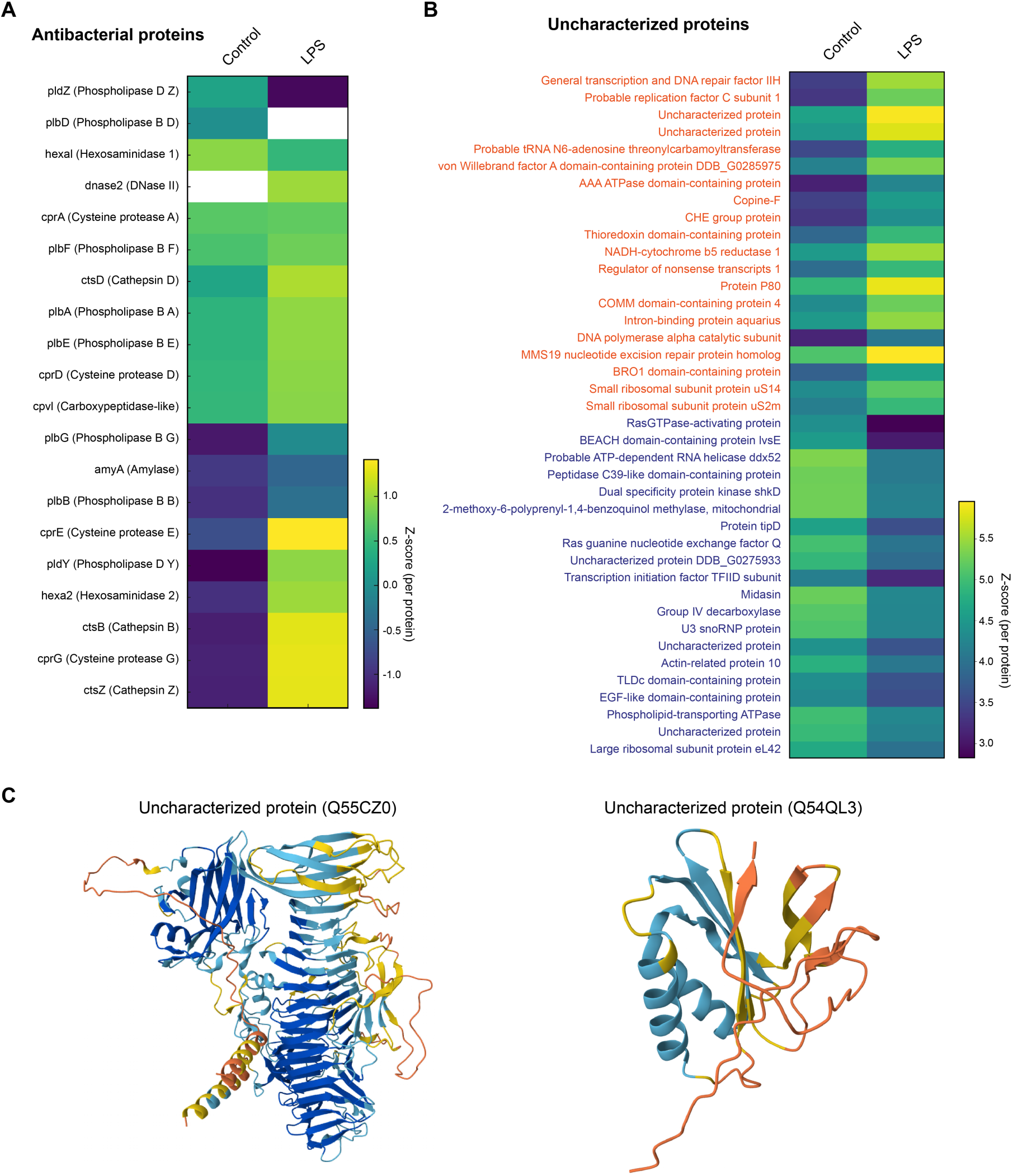
Proteomic signature of antibacterial and uncharacterized proteins in DNA extracellular traps (ETs). Heatmaps show differentially abundant proteins identified in the DNase-treated ET fraction of vegetative *Dictyostelium discoideum* cells under control and *K. pneumoniae* LPS-stimulated conditions. Proteins are grouped into antibacterial (left) and uncharacterized (right) categories. Values represent Z-score–normalized log₂-transformed protein abundances. Antibacterial proteins include phospholipases, cysteine proteases, and nucleases, whereas the uncharacterized group comprises proteins with unknown function but that are consistently enriched in ETs. The bottom panels show AlphaFold-predicted structures of two representative uncharacterized proteins (Q55CZ0 and Q54QL3), revealing well-defined folds suggestive of functional roles within ETs.

Proteomic analysis revealed a set of consistently enriched proteins with no functional annotation (Figure 5B). Differential abundance analysis identified several uncharacterized proteins selectively enriched in LPS-induced ETs. Structural prediction using AlphaFold indicated well-defined folds with features compatible with DNA interaction or regulatory functions, including conserved β-sheet architectures and putative binding grooves (Figure 5B).

Gene Ontology enrichment analysis indicated that, while the core composition of ETs is conserved, LPS stimulation subtly reshapes their functional profile. Both the control and LPS-induced ET proteomes were enriched for processes related to cytoplasmic translation, actin filament organization, and responses to bacteria (Figure 5A). However, LPS-induced ETs showed selective enrichment of nuclear protein import-related functions, whereas control ETs were enriched in cell-substrate adhesion pathways, indicating stimulus-dependent modulation of ET composition.

Together, these analyses define DNA extracellular traps formed by vegetative *D. discoideum* as mitochondrial-derived, protein-rich structures enriched in DNA-associated and antimicrobial components, with a conserved core composition and stimulus-dependent variability.

### Extracellular DNA traps are formed in response to diverse bacterial encounters

Although extracellular trap (ET) formation has been classically described in neutrophils as a response to pathogenic challenge, it remains unclear whether ET deployment by vegetative *D. discoideum* is similarly restricted to interactions with virulent bacteria. To determine whether ET formation is pathogen-specific or occurs more broadly during bacterial encounters, we examined ET deployment across distinct bacterial contexts using live-cell imaging and ultrastructural analysis.

Live-cell imaging with SYTOX Orange revealed robust formation of extracellular DNA structures upon bacterial exposure, consistent with ETs (Figure 6A; Movie S2). Interactions with virulent bacteria led to pronounced accumulation of extracellular DNA over time. DNase treatment abolished the extracellular SYTOX signal, confirming the DNA-based nature of these structures (Figure S3A). Notably, DNase-mediated disruption of ETs was accompanied by a marked late-stage increase in bacterial fluorescence (Figure S3B), indicating enhanced bacterial persistence or proliferation when ET integrity was compromised.

**Figure 6.**
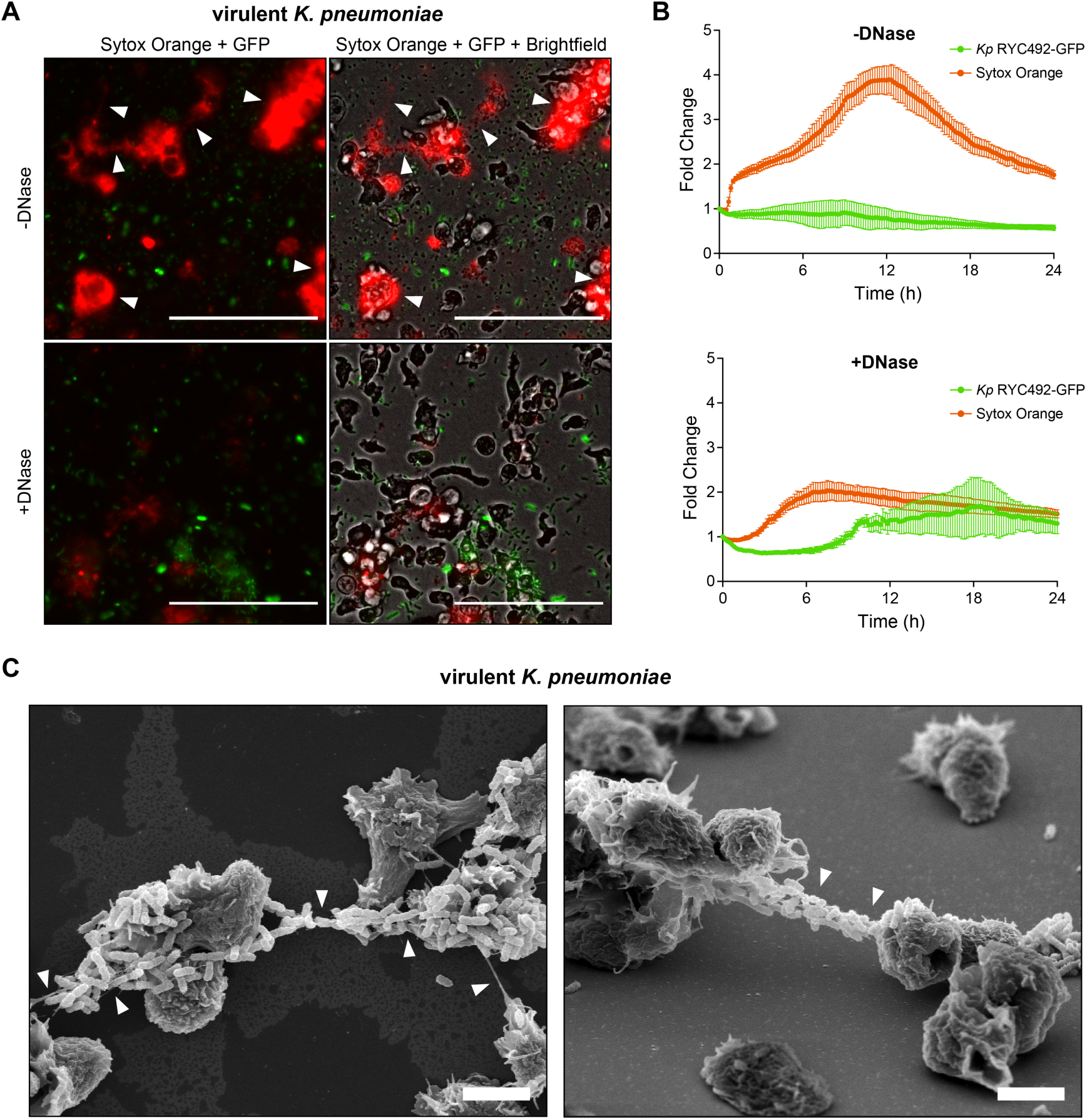
DNA ET formation in response to virulent *K. pneumoniae*. (A) Fluorescence microscopy of AX2 cells exposed to *Kp* RYC492-GFP, with and without DNase treatment. Arrowheads show ETs stained with Sytox Orange. Scale bars: 30 µm. (B) ET production and total bacterial counts were monitored by measuring Sytox Orange and GFP fluorescence over time, respectively. Fluorescence values were quantified from time-lapse fluorescence microscopy images and expressed as fold change relative to the initial time point (t = 0). (C) Ultrastructural visualization of ETs by SEM 6 h post-stimulation with Kp RYC492. Arrowheads indicate fibrillar structures projecting from *D. discoideum* cells, consistent with extracellular DNA strands. Scale bars: 5 µm.

To determine whether ET formation was restricted to pathogenic interactions, we next examined amoebal responses to bacteria commonly used as food sources in laboratory cultures and to an avirulent *E. coli* strain. In all cases, clear ET formation was observed in response to edible *K. pneumoniae* (Movie S3) and *E. coli* (Movie S4), with extracellular SYTOX-positive structures readily detected over time (Figure 7). These results demonstrate that ET deployment by vegetative *D. discoideum* occurs during encounters with both virulent and non-pathogenic bacteria.

**Figure 7.**
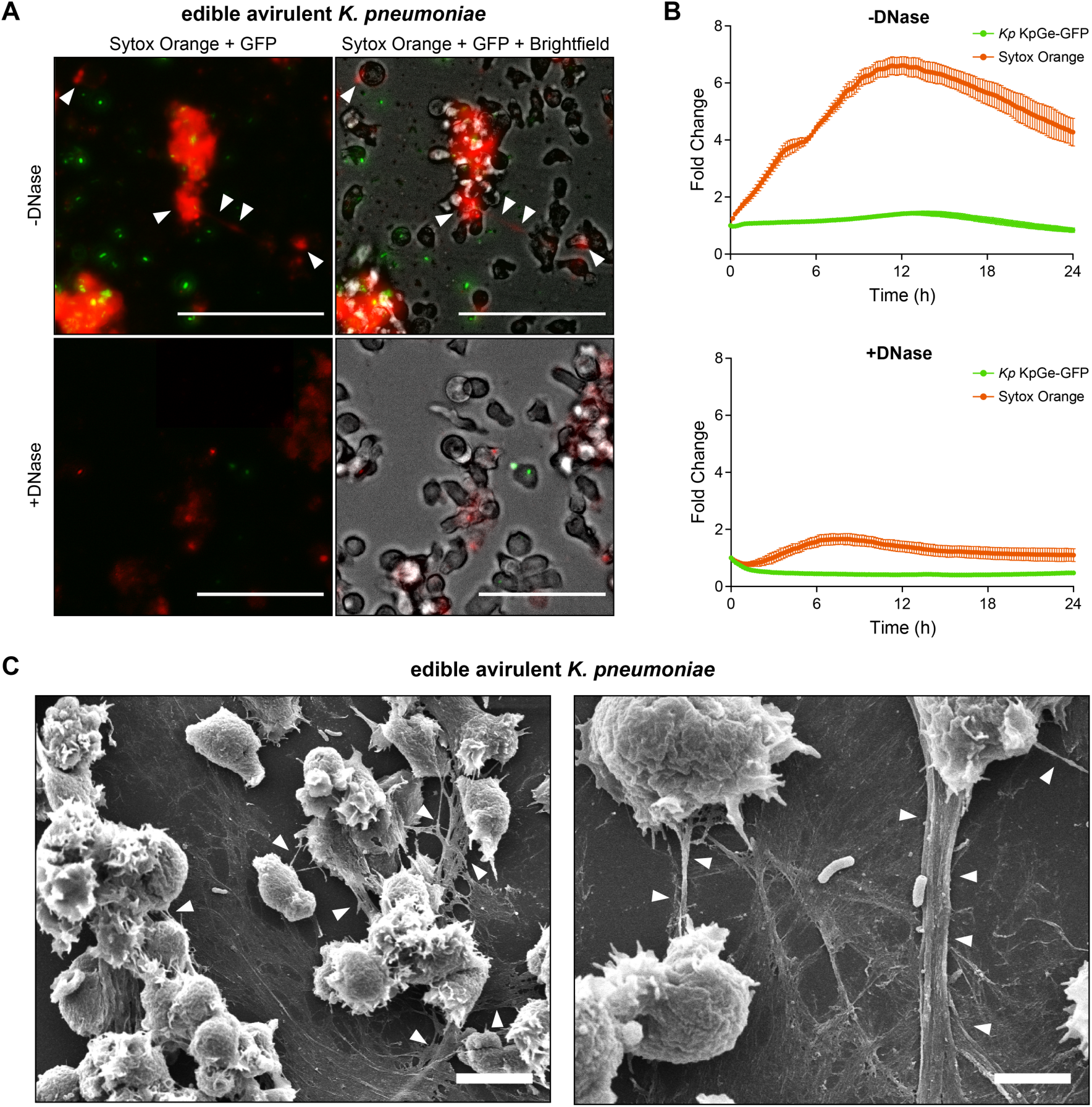
DNA extracellular traps are formed in response to edible *Klebsiella pneumoniae* KpGe. (A) Fluorescence microscopy of AX2 cells exposed to KpGe-GFP, with and without DNase treatment. Arrowheads show ETs stained with Sytox Orange. Scale bars: 30 µm. (B) ET production and total bacterial counts were monitored by measuring Sytox Orange and GFP fluorescence over time, respectively. Fluorescence values were quantified from time-lapse fluorescence microscopy images and expressed as fold change relative to the initial time point (t = 0). (C) Ultrastructural visualization of ETs by SEM 4 h post-stimulation with KpGe. Arrowheads indicate fibrillar structures projecting from D. discoideum cells, consistent with extracellular DNA strands. Scale bars: left, 10 µm; right, 5 µm.

Time-lapse imaging revealed that bacteria interacting with ETs frequently became transiently immobilized within extracellular DNA structures before eventual release or ingestion. In the case of edible bacteria, ET-associated microbes often remained extracellular for extended periods before phagocytosis. Scanning electron microscopy provided complementary structural evidence, revealing extracellular filamentous material connecting amoebae and bacteria under all bacterial exposure conditions examined. These filamentous structures were consistent with DNA-based networks observed by fluorescence microscopy and confirmed that ETs represent bona fide extracellular matrices rather than imaging artifacts.

Together, these observations demonstrate that vegetative *D. discoideum* deploys extracellular DNA traps during interactions with a broad spectrum of bacterial species, and that ET integrity limits bacterial expansion during these encounters.

## Discussion

DNA extracellular trap formation has traditionally been conceptualized as a specialized immune response, first described in vertebrate neutrophils as filamentous DNA-protein structures that immobilize microbes ^21,24^. Subsequent work has expanded this view, revealing that ETosis is not restricted to neutrophils nor is it invariably associated with cell death; rather, it represents a broader and more versatile biological strategy ^21,25^. Here, we show that vegetative cells of the social amoeba *D. discoideum* deploy extracellular DNA traps as a regulated, non-cytotoxic response to bacterial encounters. ET formation in vegetative amoebae occurs through a vital ETosis mechanism that preserves membrane integrity and feeding capacity, is selectively induced by specific bacterial cues, and involves the extracellular release of mitochondrial DNA associated with a diverse repertoire of antimicrobial proteins. Together, these findings reposition ETosis as a cellular strategy operating at the interface of microbial containment and nutrition, rather than a response restricted to canonical immune contexts.

A defining feature of ETosis in vegetative *D. discoideum* is its non-lytic nature. Functional viability assays, membrane integrity measurements, and ultrastructural analyses consistently demonstrated that ET formation occurs without loss of plasma membrane integrity or feeding competence. This vital mode of ETosis contrasts with the lytic NETosis frequently observed in neutrophils, yet closely resembles non-lytic forms of ETosis described in other immune cell types, including eosinophils and neutrophils that undergo mitochondrial DNA release under specific activation conditions. The selective enrichment of mitochondrial, but not nuclear, DNA within extracellular traps further supports a coordinated extrusion mechanism rather than passive DNA leakage associated with cytotoxic stress, consistent with previous observations in amoeboid sentinel cells and mammalian immune systems ^26–28^.

ET formation in vegetative amoebae also exhibited high stimulus specificity. While lipopolysaccharide (LPS) robustly induced ET formation, classical neutrophil NET inducers such as phorbol esters and flagellin failed to elicit comparable responses. In vertebrate immune cells, these stimuli activate protein kinase C-dependent and Toll-like receptor-mediated signaling pathways that drive NET release ^21,29^. Their inability to trigger ETosis in *D. discoideum* suggests fundamental differences in upstream sensing and signaling architecture, consistent with the notion that innate immune signaling pathways have undergone extensive evolutionary rewiring despite conservation of downstream effector strategies ^30,31^. While our data demonstrate clear stimulus specificity in ET induction, the upstream sensing mechanisms and signaling pathways linking bacterial cues to mitochondrial DNA extrusion remain unresolved. Future genetic and biochemical dissection of pattern recognition receptors and downstream signaling nodes will be required to define how microbial signals are transduced into ET deployment in amoebae.

Molecular characterization of amoeboid ETs further revealed that these structures are not inert DNA scaffolds but protein-rich extracellular assemblies. Proteomic analyses showed that ETs are enriched in mitochondrial proteins, DNA-associated factors, and a broad repertoire of enzymes with known or predicted antimicrobial activity, including phospholipases, proteases, nucleases, and glycosidases. Many of these effectors have previously been implicated in bacterial killing, digestion, or clearance in *D. discoideum* ^22,23,32,33^, and their extracellular enrichment within ETs parallels observations made in vertebrate NETs, where antimicrobial proteins contribute to microbial damage and containment ^21,25,34^. In addition, the presence of multiple uncharacterized proteins selectively enriched in ETs points to previously unexplored molecular components of extracellular trap biology.

An important insight emerging from this study is that ET formation in vegetative *D. discoideum* is not restricted to interactions with virulent or pathogenic bacteria. Edible bacteria commonly used as laboratory food sources, as well as attenuated bacterial strains, also elicited robust ET deployment. Live-cell imaging revealed that, under these conditions, bacteria frequently became transiently immobilized within extracellular DNA networks without immediate killing or phagocytosis. This behavior contrasts with the classical immunocentric view of extracellular traps as terminal antimicrobial weapons and instead indicates that ETs can function in contexts where microbial survival is temporarily tolerated.

Consistent with these observations, the retention of bacteria within ETs for extended periods suggests a role in spatial organization and microbial management rather than exclusively in microbial elimination. In the case of edible bacteria, ET-associated microbes were often observed to remain confined outside the cell before eventual release or ingestion. Comparable extracellular holding strategies have been described in contexts of protist grazing and environmental predation ^35–38^, supporting a broader ecological function for ETosis. Although live-cell imaging and DNase disruption assays support a role for extracellular traps in microbial containment, the precise impact of ET formation on long-term bacterial fitness, digestion efficiency, and nutrient acquisition remains to be quantitatively resolved.

High-resolution fluorescence microscopy and scanning electron microscopy provided direct evidence that extracellular traps formed by vegetative *D. discoideum* physically capture bacteria. Extracellular DNA filaments were repeatedly observed entrapping bacterial cells in close association with amoebae, while ultrastructural analyses revealed filamentous extracellular matrices connecting amoebae and bacteria. This physical immobilization closely resembles extracellular DNA-mediated trapping, as described in neutrophils and other immune cells, in which extracellular traps limit microbial dissemination and facilitate downstream clearance ^39,40^.

The functional relevance of ET-mediated containment was further supported by DNase-mediated disruption experiments. Enzymatic degradation of extracellular DNA was associated with increased bacterial signal over time, particularly during interactions involving virulent bacteria, consistent with reduced containment. Similar containment functions have been attributed to neutrophil extracellular traps in vertebrates, where physical trapping restricts microbial spread even in the absence of immediate killing ^39,40^.

From an evolutionary perspective, these findings align with a growing body of evidence proposing that many traits traditionally interpreted as virulence or immune factors originated as ecological adaptations shaped by interactions with environmental hosts ^36,41–44^. Protist predation has been identified as a major selective force driving the evolution of bacterial traits that subsequently contribute to animal pathogenesis, including resistance to phagocytosis and stress tolerance ^36–38^. Within this framework, ETosis may have emerged as an ancestral strategy employed by phagocytic eukaryotes to immobilize, manage, and process bacteria in microbe-rich environments.

Recent discoveries of extracellular DNA trap formation in early-diverging metazoans further reinforce the deep evolutionary roots of this trait ^20,45^. Cells capable of deploying ET-like structures have been identified in non-bilaterian animals, supporting the view that extracellular DNA-based microbial containment predates the evolution of complex immune systems ^46^. While comparative observations across protists and early-diverging metazoans support an ancient origin of extracellular DNA-based microbial containment, direct phylogenetic and cross-species functional analyses will be necessary to formally reconstruct the evolutionary trajectory linking nutritional predation mechanisms to modern immune extracellular traps. In a manner reminiscent of phagocytosis, which is thought to have originated as a nutritional strategy in early eukaryotes and was later co-opted for immune defense functions ^47–49^, our findings suggest that extracellular trap formation may have followed a similar evolutionary trajectory. By demonstrating that unicellular eukaryotes deploy ETs not only for defense but also to manage nutritional interactions, this work supports a unifying view of ETosis as an evolutionarily ancient and adaptable strategy for microbial management within eukaryotic cells ^30^.

In summary, this study establishes that extracellular trap formation in vegetative *D. discoideum* is a regulated, multifunctional response to bacterial encounters, integrating physical containment, biochemical activity, and spatial organization. By demonstrating that ETosis operates in a unicellular phagocyte at the intersection of microbial management and nutrition, our findings challenge a strictly immunocentric view of extracellular traps. Instead, they support a model in which ETosis represents an evolutionarily ancient strategy for handling microbes that was later adapted and refined in multicellular immune systems.

Understanding extracellular traps through this broader ecological and evolutionary lens provides a framework for reinterpreting their roles across diverse biological contexts and for exploring how fundamental cellular strategies are repurposed during the evolution of immunity.

## Material and Methods

### Cell Culture

Frozen aliquots of axenic *D. discoideum* AX2 (or AX4) were thawed and transferred to Petri dishes containing HL5 medium^50^. Cells were maintained at 23°C and grown to 80–90% confluence; at this point, they were split into new plates at varying densities, as required by the experiment.

### Microplate Assays for Extracellular Trap (ET) Quantification

To evaluate ET formation, 96-well black plates were used. *D. discoideum* AX2 and AX4 cells were seeded at 2 × 10⁵ cells/200 μL in Soerensen buffer, supplemented with 5 μM Sytox Green, a non-cell-permeable DNA-binding dye, and incubated for 20 min at 23°C. Cells were then stimulated with 2.5, 5, or 10 µg/mL LPS from *K. pneumoniae*.

Fluorescence was measured every 20 min for 4 hours using a Tecan Infinite 200Pro microplate reader (excitation: 480/490 nm, emission: 520 nm). After the assay, cells were examined under a fluorescence microscope to confirm viability and to ensure that fluorescence originated primarily from extracellular DNA.

### Extracellular Trap Visualization and Live-Cell Imaging

For static imaging, *D. discoideum* cells were suspended in Soerensen buffer containing Sytox Green and seeded at 3 × 10⁵ cells per well in 300 µL of medium in 24-well plates. After 20 min, cells were stimulated with 10 µg/mL LPS. Images were captured every 10 min at 20× magnification using the Lionheart FX automated fluorescence microscope (in the brightfield and green fluorescence channels). Additional imaging was performed using a confocal microscope (Zeiss model LSM 700). To assess the DNA composition of ETs, cells were prepared as described above and stimulated with 10 µg/mL LPS for 4 hours. Individual ET structures were identified in the brightfield and green fluorescence channels, and their coordinates (x, y) were recorded. Turbo DNase (10 U) and 10× DNase buffer were carefully added to wells to avoid cell disruption. The plate was incubated at 23°C for 40 min, after which the same ET structures were re-imaged using identical GFP channel settings.

### Plaque Viability Assay

A suspension of *D. discoideum* AX2 or AX4 cells (1×10⁵ cells/mL in HL5 medium) was diluted to 1×10⁴ cells/mL, and 5 mL aliquots were treated with one of the following: 10 µg/mL lipopolysaccharide (LPS) from *Klebsiella pneumoniae* (Sigma-Aldrich, cat. L4268) or 10 µg/mL G418 (Gibco, cat. 10131-035). Cells were incubated for 6 hours at 23°C with agitation (150 rpm).

To assess viability, *K. pneumoniae* KpGe (strain DBS0351098) was cultured overnight, washed, and resuspended in 1X Sorensen buffer. A 960 µL bacterial suspension was mixed with 20 µL (200 cells) from each experimental condition and plated on SM agar. After 3 days of incubation at 23°C, phagocytic plaques were counted as a proxy for viable cells.

### Flow Cytometry Viability Assay

A. *D. discoideum* AX2 cells were washed and resuspended in filtered (0.22 µm) 1X Sorensen buffer at 5×10⁵ cells/mL. Aliquots (1 mL) were diluted to 2.5×10⁵ cells/mL and treated with 10 µg/mL *K. pneumoniae* LPS, or 10% ethanol (viability control).

After 6 hours of incubation (23°C, 150 rpm), 1 mL of each sample was stained with propidium iodide (0.1 µg/mL final concentration) and analyzed on a BD FACSCanto II flow cytometer (excitation: 488 nm; emission: 655 nm). A total of 10,000 events were recorded per sample. Data were processed using FACSDiva software.

### Live Imaging of Bacteria-Induced ET Formation in *D. discoideum*

For bacteria-exposed ET formation assays, *D. discoideum* AX4 cells were seeded at 1 × 10⁶ cells/mL in 24-well plates, centrifuged at 2400 rpm for 10 min, and incubated at 23°C for 24 hours to allow adhesion. After three washes with Soerensen buffer, bacterial suspensions (OD₆₀₀ = 1 × 10⁷ cells/mL, MOI = 10) were added to wells and centrifuged at 2400 rpm for 30 min to facilitate interaction.

For ET detection, SYTOX Orange (250 nM) was used to stain extracellular DNA. Turbo DNase (20 U) was added for degradation experiments. Live imaging was performed using the Lionheart FX, capturing images every 10 minutes for 24 hours in brightfield and fluorescence channels. Fluorescence intensity was analyzed using the Time Series Analyzer V3 plugin in ImageJ. Total fluorescence intensity was measured for each image over time, and the resulting data tables were exported for further analysis. Fluorescence intensity values were then normalized by calculating the relative change in fluorescence intensity over time.

### Isolation of ET Fractions

To determine the DNA origin of ETs, *D. discoideum* cells were seeded at 4 × 10⁶ cells per mL in 2 mL of Sørensen buffer in 12-well plates and stimulated with 10 µg/mL LPS for 4 hours. Control wells received only buffers. After induction, three fractions were collected: (1) Secretome: the upper supernatant layer, transferred to a clean tube (not used for qPCR). (2) ET Fraction: the middle layer was collected carefully and centrifuged at 5000 rpm for 3 min to remove debris. This fraction was stored at 20°C for qPCR. (3) Cell Fraction: the remaining adhered cells were collected and subjected to HotSHOT DNA extraction ^51^. qPCR was performed using SYBR Green Master Mix, targeting nuclear (*H3a*) and mitochondrial (*rnlA*) genes, with three biological replicates and three technical replicates per condition. Data normalization was performed using Pfaffl’s method, and unpaired t-tests were used to compare conditions. Following induction, the ET fraction was carefully collected. The supernatant was then precipitated with 100 mM LiCl and 2 volumes of ethanol at 4°C, followed by overnight incubation at -80°C. The next day, precipitated nucleic acids were pelleted by centrifugation at 17,000 rpm for 15 minutes at 4°C. The pellet was then resuspended in 25 µL of nuclease-free water. Samples were treated with Turbo DNase in the presence of 10× DNase buffer before being loaded onto a 1.5% agarose gel in TAE buffer for electrophoresis.

### Global Proteomic Profiling of ETs

For mass spectrometry analysis, the ET fraction and secretome were processed. *D. discoideum* cells were seeded at 4 × 10⁶ cells in 2 mL Soerensen buffer in 12-well plates and stimulated with 10 µg/mL LPS from *K. pneumoniae* for 4 hours. Each condition was prepared in quadruplicate. After separation, the ET fraction was treated with Turbo DNase (5 U) for 40 min; 400 µL was then collected and supplemented with 15 mM EDTA to inactivate the DNase. This fraction was centrifuged at 5000 rpm for 3 min to remove debris and stored for mass spectrometry analysis. Before MS analysis, an SDS-PAGE silver stain was performed to confirm the presence of proteins.

Proteins were extracted using chloroform/methanol precipitation and digested with trypsin at a 1:50 protease-to-protein ratio. Peptides were cleaned using Sep-Pak C18 Spin Columns (Waters), dried, and analyzed using nanoUHPLC nanoElute coupled to a timsTOF Pro mass spectrometer (Bruker Daltonics). Data collection employed TimsControl 2.0 with 10 PASEF cycles over a 100-1700 m/z mass range. Protein abundance data were analyzed using MSFragger v3.5 through Fragpipe v18.0 ^52^. Proteins were identified using the D. discoideum Uniprot proteome with 1% FDR filtering.

Data were processed using R with specialized proteomics packages. Proteins with spectral counts below three were excluded. Data were log₂-transformed and median-centered, and log-fold changes between conditions were computed. Statistical significance was assessed using two-sample t-tests with Benjamini-Hochberg correction ^53^.

Gene Ontology (GO) enrichment was performed using DAVID (v2023q4) ^54^, with results reported using Benjamini-adjusted p-values. Structural homology searches for uncharacterized proteins were conducted using Foldseek ^55^, with AlphaFold used to predict *D. discoideum* protein structures ^56,57^.

### Scanning electron microscopy

Scanning electron microscopy (SEM) was used to visualize the ultrastructure of extracellular DNA traps (ETs) formed by vegetative *D. discoideum* cells following bacterial or LPS stimulation. Amoebae were allowed to adhere to suitable substrates and were stimulated under ET-inducing conditions; after which, samples were fixed at defined time points using aldehyde-based fixation to preserve cellular and extracellular structures. Fixed samples were subsequently dehydrated through a graded ethanol series, critical-point-dried to preserve filamentous ET architecture, and sputter-coated with a conductive metal layer.

SEM imaging was performed using a field-emission scanning electron microscope (Zeiss EVO MA10, Germany) under high-vacuum conditions, allowing high-resolution visualization of cell morphology, extracellular DNA fibers, and bacteria-associated ET networks. Images were acquired at multiple magnifications to resolve both global ET organization and fine filamentous details characteristic of extracellular DNA-based structures.

## Supporting Information

**Figure S1. Differential induction of extracellular traps by LPS of distinct bacterial origin.** (A) Sytox Green fluorescence over time in AX2 cells stimulated with 10 µg/mL lipopolysaccharide (LPS) from *K. pneumoniae* (*Kp* LPS), *E. coli* (*Ec* LPS), or *P. aeruginosa* (*Pa* LPS). (B) At 4 h post-stimulation, fold change in fluorescence intensity relative to untreated controls is shown for the three concentrations tested of *Ec* LPS and *Pa* LPS. For *Pa* LPS, a significant increase in ET formation is observed only at the highest concentration. For *Ec* LPS, no statistically significant induction was detected, although a trend toward increased fluorescence was observed. Data are presented as mean ± SD (n = 3 biological replicates). Statistical significance was assessed using one-way ANOVA with Dunnett’s multiple comparisons test (*p < 0.05). (C) Representative fluorescence microscopy images illustrating differences in extracellular trap density 4 h post-stimulation with 10 µg/mL LPS. Scale bar, 50 µm.

**Figure S2. PMA and flagellin do not induce extracellular trap formation in vegetative** (A) ***D. discoideum*.** AX2 cells were stimulated with increasing concentrations of phorbol 12-myristate 13-acetate (PMA; 50, 500, and 1000 nM) (A) or bacterial flagellin (100, 200, and 300 ng/mL) (B). Extracellular trap production was quantified by measuring Sytox Green fluorescence over time. At 4 h post-stimulation, fluorescence intensity fold change in fluorescence intensity relative to untreated controls confirms the absence of significant ET induction under all conditions tested. Data are presented as mean ± SD (n = 3 biological replicates). Statistical significance was assessed using one-way ANOVA with Dunnett’s multiple comparisons test.

**Figure S3. DNase treatment disrupts extracellular traps during bacterial encounters.** (A) Fluorescence microscopy of AX2 cells exposed to *E. coli* B/r-GFP, in the presence or absence of DNase treatment. Arrowheads indicate extracellular traps stained with Sytox Orange. Scale bars, 30 µm. (B) Extracellular trap formation and bacterial abundance were quantified over time by measuring Sytox Orange and GFP fluorescence, respectively. Fluorescence values were extracted from time-lapse microscopy images and expressed as fold change relative to the initial time point (t = 0).

**Movie S1. Confocal z-stack imaging of AX2 Dictyostelium discoideum cells stimulated with lipopolysaccharide (LPS).** A cluster of vegetative amoebae is shown surrounded by an extensive filamentous extracellular network stained with Sytox Green, corresponding to extracellular DNA traps. The three-dimensional reconstruction highlights the spatial organization of extracellular trap structures relative to the cell aggregate. Scale bar: ∼10 µm.

**Movie S2. Time-lapse fluorescence microscopy showing the interaction between vegetative AX2 *D. discoideum* cells and the virulent *K. pneumoniae* strain RYC492 expressing green fluorescence.** Extracellular DNA is visualized using Sytox Orange, revealing the dynamic formation and remodeling of extracellular traps during bacterial encounters. The movie illustrates the spatiotemporal deployment of DNA-based filamentous networks in close association with bacterial cells. Scale bar: ∼10 µm.

**Movie S3. Time-lapse fluorescence microscopy showing the interaction between vegetative *D. discoideum* cells and the edible, avirulent *K. pneumoniae* KpGe strain expressing green fluorescence.** Extracellular DNA is visualized using Sytox Orange, revealing the dynamic formation of extracellular traps during encounters with non-virulent bacteria. The movie highlights the deployment of DNA-based filamentous networks in response to a bacterial food source. Scale bar: ∼10 µm.

**Movie S4. Time-lapse fluorescence microscopy showing the interaction between vegetative *D. discoideum* cells and the edible, avirulent *E. coli* B/r strain expressing green fluorescence.** Extracellular DNA is visualized using Sytox Orange, revealing the dynamic formation of extracellular traps during encounters with a bacterial food source. The movie illustrates the deployment of filamentous extracellular DNA networks in association with non-pathogenic bacteria. Scale bar: ∼10 µm.

## Author contribution

Antonia Ramos-Guzmán contributed to conceptualization, investigation, methodology, formal analysis, and writing of the original draft, particularly performing LPS experiments and proteomic analyses. Paulina Aguilera-Cortés contributed to the investigation and methodology related to PAMPs experiments, as well as writing the original draft and reviewing and editing the manuscript. Tabata Soto conducted electron microscopy experiments. Boris Riveros performed flow cytometry experiments. Mauricio Hernández carried out formal analysis and data curation of proteomics datasets. Ian Pérez contributed to conceptualization, experimental design and investigation in *Dictyostelium discoideum*, and writing of the original draft. Diego Rojas was responsible for sample preparation for microscopy, particularly scanning electron microscopy (SEM). Sebastián Farías performed live-cell imaging experiments with bacteria. Viviana Barros contributed to the investigation of PAMPs, including LPS and other stimuli. Andrés E. Marcoleta and Francisco P. Chávez conceived the study, supervised the research, contributed to writing (original draft, review, and editing), and secured funding.

## Acknowledgment

This work was funded by Fondecyt Grants 1262424 (FCH). Thank Nicole Molina for her excellent technician skills in preparing *D. discoideum* cultures.

